# Time dependent response of daunorubicin on cytotoxicity, cell cycle and DNA repair in acute lymphoblastic leukaemia

**DOI:** 10.1101/346031

**Authors:** Hussain Mubarak Al-Aamri, Heng Ku, Helen R Irving, Joseph Tucci, Terri Meehan-Andrews, Christopher Bradley

**Author notes:** Corresponding author: (HRI).

## Abstract

Daunorubicin is commonly used in the treatment of acute lymphoblastic leukaemia (ALL). Various mechanisms of action for daunorubicin have been proposed and its action is likely to be multi-modal. The aim of this study was to explore the kinetics of double strand break (DSB) formation of three ALL cell lines following exposure to daunorubicin and to investigate the effects of daunorubicin on the cell cycle and the protein kinases involved in specific checkpoints following DNA damage and recovery periods. Three ALL cell lines CCRF-CEM and MOLT-4 derived from T lymphocytes and SUP-B15 derived from B lymphocytes were examined following 4 hours treatment with daunorubicin chemotherapy and varying recovery periods. Daunorubicin induced different degrees of toxicity in all cell lines and consistently generated reactive oxygen species. Daunorubicin was more potent at inducing DSB in MOLT-4 and CCRF-CEM cell lines while SUP-B15 cells showed delays in DSB repair and significantly more resistance to daunorubicin compared to the other cell lines as measured by γH2AX assay. Daunorubicin also causes cell cycle arrest in all three cell lines at different checkpoints at different times. These effects were not due to mutations in Ataxia–telangiectasia mutated (ATM) as sequencing revealed none in any of the three cell lines. However, p53 was phosphorylated at serine 15 only in CCRF-CEM and MOLT-4 but not in SUP-B15 cells. The lack of active p53 may be correlated to the increase of SOD2 in SUP-B15 cells. The delay in DSB repair and lower sensitivity to daunorubicin seen in the B lymphocyte derived SUP-B15 cells could be due to loss of function of p53 thus causing variations in the DNA repair pathways.

## Introduction

Daunorubicin is an anthracycline antibiotic that is widely used in treating acute leukaemia, lymphoma and multiple myeloma [1]. Proposed mechanisms of anthracycline action have included: inhibition of synthesis of macromolecules through intercalation of daunorubicin into DNA strands [2, 3], interaction with molecular oxygen to produce reactive oxygen species (ROS), topoisomerase II (TOPO2) inhibition and the formation of DNA adducts [4]. There is good evidence for all these pathways and the mechanism of action of the anthracyclines is likely to be multi-modal. The type of toxic lesions that generally results from daunorubicin treatment are DNA double strand breaks (DSB). The occurrence of DSB activates PI3K-like kinases such as Ataxia–telangiectasia mutated (ATM) [5]. ATM exists as an inactive dimer and undergoes autophosphorylation and monomerisation in response to DNA DSB [6]. Activated ATM phosphorylates H2AX at Ser139 residues of the carboxyl terminus to form γH2AX around the DNA-DSB. A large number of γH2AX molecules form around the DSB to create a focus point where various DNA repair and checkpoint proteins accumulate that facilitate DNA-DSB repair [7]. In response to DNA DSB, ATM initiates repair by either non-homologous end joining (NHEJ) or homologous recombination (HR) though the factors controlling which pathway is chosen are not well understood [8]. A common outcome of both pathways is phosphorylation of the tumour suppressor gene, p53, which plays a pivotal role in the cellular response to damage as p53 regulates numerous cellular responses, including cell cycle arrest and apoptosis as well as upregulation of anti-oxidant proteins such as manganese-containing superoxide dismutase (SOD2 or MnSOD) [9].

Phosphorylation of p53 is an essential factor for the activation of key cell cycle checkpoints that leads to a delayed cell cycle progression, resulting in a reversible arrest at the G1/S cell cycle checkpoint [10] and is also involved in the arrest of the G2/M checkpoint [11]. The activation of these checkpoints allows more time for DNA repair mechanisms to be initiated to maintain genomic integrity[10].

Increased levels of ROS following daunorubicin treatment can directly activate ATM in vitro [12]. It is proposed that ROS activates ATM by promoting the formation of disulphide bridges, and thus stabilising the ATM dimer, rather than forming a monomer as follows activation by DSBs. Since activated ATM remains as a dimer, ATM may engage a different set of substrates and thus different cellular responses. While there is subsequent downstream activation of p53 and other proteins activated by DSB, the other downstream targets of ATM activated by ROS are thought to differ substantially [12]. This could have potential effects on cell cycle arrest and the initiation of apoptosis as well as cellular redox homeostasis.

The process of lymphoid tumourigenesis often involves alterations to the ATM gene resulting in ATM deficient cells which are more sensitive to oxidative stress and are likely to undergo altered DNA repair and apoptotic pathways. We have chosen to limit our study to acute lymphoblastic leukaemia (ALL) lines as daunorubicin is widely used in the treatment of this leukaemia [13]. Little is known about the ATM sequence in ALL cell lines used in medical research. One of the aims of this study was to explore potential functional mutations in ATM that may affect how cells handle chemotherapy treatment. To this end, the ATM coding sequences in T-lymphoblast derived CCRF-CEM and MOLT-4 cells and B-lymphoblast derived SUP-B15 cells were analysed through Sanger sequencing.

Following daunorubicin treatment, activation of ATM occurs via different mechanisms. Firstly, ATM activation by DSB involves autophosphorylation at Ser1981 and monomerization of the native dimer, and subsequent phosphorylation of H2AX to form γH2AX [14]. We monitored DSB formation by measuring the formation of γH2AX following treatment of several ALL cell lines with daunorubicin. As the repair process is dynamic and may involve sequential involvement of different repair pathways we analysed DSB over a time course. Secondly, as ROS production may activate ATM, we measured ROS production following exposure of ALL cells to daunorubicin. The impact γH2AX and ROS levels have on ATM function and cell survival were analysed.

## Materials and methods

### Cell lines

Two T-lymphoblastic leukaemia cell lines, CCRF-CEM and MOLT-4, and a Blymphoblastic leukaemia cell line, SUP-B15, were obtained from the American Type Cell Culture Collection (ATCC). Cells were stored frozen in liquid nitrogen in cryovials until use. The CCRF-CEM and MOLT-4 cells were cultured in RPMI-1640 (GibcoTM-Life Technology, NY, USA), while the SUP-B15 cells were cultured in IMDM (GibcoTM-Life Technology, NY, USA) supplemented with 10 % foetal calf serum (FCS) and 5 mM glutamine. Cells were incubated at 37°C in aerobic atmosphere containing 5% CO_2_. Cells were used before 10 passages.

### Daunorubicin treatment and recovery

Daunorubicin (Sigma-Aldrich NSW, Australia) was prepared as 5 mM stock solutions in dimethyl sulfoxide (DMSO) and stored in aliquots at −20°C. Cells were plated onto six well plates at a seeding density of 1×10^6^ cells per well. Cells were treated with 10 μM daunorubicin and incubated for 4 hours at 37°C in atmosphere of 5% CO_2_. Treatment media was removed by centrifugation at 200 g for 5 minutes and replaced with recovery media (media only) for 4, 12 or 24 hours.

### MTT (3-(4,5-dimethylthiazol-2-yl)-2-5 diphenyltetrazolium bromide) assay

After treatment, cells were washed with PBS (phosphate buffer saline) and centrifuged at 4°C at 500 g for 5 minutes. The cells were then plated into 96-well plate (3 x 10^4^ cells per well), and 0.25 mg/ml MTT (Sigma-Aldrich) was added to each well and then incubated at 37°C for 3 hours protected from light. Formazan crystals were solubilized by incubation in 10 % DMSO at room temperature for 1½ hours before reading absorbance at 570 nm using a Flex station 3 (Molecular Devices, California, USA).

### ROS flow cytometry assay

The cells were plated onto six well plates at a seeding density of 1 × 10^6^ cells per well, prior to treatment and recovery. The cells were then collected and centrifuged at 400 g for 5 minutes before resuspending in fresh media at 1 x 10^5^ cells ml^−1^. One ml of cell suspension was added to 1.5 ml microcentrifuge tubes. As a negative control 5 mM of ROS inhibitor (N-acetyl-L-cysteine) (Enzo-life sciences, NY, USA) was added at least 30 minutes prior to induction, while 500 μM pyocyanin (ROS inducer) was used as a positive control [15]. ROS detection solution (5 mM, Enzo-life sciences) was added to all tubes before incubation at 37° C for 30 minutes in the dark. Finally, the intensity of cell fluorescence was recorded by flow cytometry (Accuri^®^ C6, Flow cytometery, Ann Arbor,MI, USA) using 500 nm excitation and 600 nm emission.

### Gamma H2AX assay

Cells were plated in a six well plate at a seeding density of 2 × 10^5^ cells per well, prior to treatment and recovery. The cells were then transferred into 1.5 ml microcentrifuge tubes and centrifuged at 200 g for 5 minutes. After centrifugation, cells were washed twice with ice cold TBS (Tris buffered saline, pH 7.4) to remove traces of ethanol in samples. All the samples were kept on ice during assay procedure. After washing with TBS, cells were resuspended in 500 μl of ice cold TFX (1 x TBS, 4% FCS, 0.1% Triton-X100 made fresh for each experiment) and allowed to rehydrate for 10 minutes. Cells were centrifuged again at 200 g and supernatant was removed. Cells were resuspended in 100 μl anti-H2AX (pSer139) rabbit polyclonal IgG (ThermoFisher Scientific, Waltham, MA, USA) diluted at 1:500 in 1x TFX and incubated at room temperature for 2 hours. Cells were then washed twice with 1x TFX by centrifugation at 200 g and the supernatant discarded. Cells were resuspended in 100 μl goat anti-rabbit IgG Alexa Fluor 488 conjugate antibody (ThermoFisher Scientific, USA) diluted at 1:200 in 1x TFX for 1 hour at room temperature, protected from light. Cells were washed twice with 1x TFX and resuspended in 300 μl TFX containing 5 μg ml^−1^ propidium iodide (Sigma Aldrich) and analysed using flow cytometery. Data was analysed using CFlow Plus software (Accuri®). Log fluorescence against cell count was plotted.

### Cell cycle analysis

Fixed samples were centrifuged at 200 g and washed twice with ice-cold PBS. The cells then resuspended in staining solution containing 25 μg ml^−1^ propidium iodide and 100 μg ml^−1^ RNase A in cold PBS and incubated at 37°C for 30 minutes. All the samples were analysed by using a flow cytometer (BD Accuri C6, California, USA). A total of 10,000 events were recorded for each sample.

### Western blotting and dot array

After treatment with daunorubicin, cells were washed with PBS containing 1 mM phenymethylsulfonyl fluoride (PMSF) and centrifuged at 200 g for 5 minutes at 4°C. Pelleted cells were suspended with 100 μl lysis buffer (Abcam, VIC, Australia), 1 mM PMSF and 10 mg ml^−1^ aprotinin, and incubated on ice for 30 minutes. Samples were then centrifuged at 15,000 g for 20 minutes at 4°C and the supernatants collected. Protein concentration of each sample was determined using BSA protein assay according to the manufacturer’s protocol (BIO-RAD, NSW, Australia). Each protein sample (30 μg) was denatured in 5 μl LDS (lithium dodecyl sulfate buffer) with 200 mM DTT at 70°C for 10 minutes (Abcam). Samples were loaded onto 10% or 12% Tris Tricine SDS-PAGE gels (Abcam) and fractionated at 180 V for 30 to 70 minutes in the presence of SDS running buffer (Abcam, ab 119195). Samples were transferred to polyvinylidene fluoride (PVDF) membrane using a mini trans-blot apparatus according to the manufacture’s protocol (BIO-RAD) with 1x transfer buffer (Abcam). Following protein transfer, the PVDF membrane was blocked with 5% non-fat milk in TBS-T (20 mM Tris, pH 8.0, 150 mM NaCl, 0.1% (v/v) Tween 20) overnight. The PVDF membrane was washed three times with TBS-T each for 5 minutes and then incubated in either anti-SOD2 (1:1000), anti-p53 (1:1000) or anti-beta tubulin (1:5000) antibodies (Cell Signalling Technology, MA, USA) diluted in blocking buffer for 4 hours at 4°C with gentle agitation. The membrane was then washed with in TBS-T three times, before adding goat anti-rabbit IgG secondary antibody (1:10,000) (Cell Signalling Technology) and incubating for 4 hours at room temperature. The PVDF was then washed with TBS-T and incubated with ChemiFast Chemiluminescence substrate (BIO-RAD). Chemiluminescence was measured in a G BOX (Syngene, Cambridge, UK) and the immunodensity of the bands was measured using gene tool Syngene software.

The impact of treatment on proteins involved in cell cycle arrest was explored using a commercially available human apoptosis array kit (Abcam, UK) according to the manufacturer’s instructions. After treatment with daunorubicin, cells were incubated for a further 12 hours in recovery media. Cells were removed from plates and processed according to the manufacturer’s instructions Sample lysate protein concentration was determined with BioRad DC Protein Assay Kit II before blotting and immunoprobing and detection of chemiluminescence signal as described above.

## ATM Sequencing

RNA was isolated from 5 x 10^6^ of CCRF-CEM, MOLT-4 or SUP-B15 cells using SV total RNA isolation kit (Promega, VIC, Australia). Reverse transcription was performed using the ImProm-IITM Reverse transcription system (Promega, VIC, Australia) kit. 1 μg of RNA samples were mixed with 0.5 μg of oligo dT, 0.5 μg of random primers and nuclease-free water. The reverse transcription mix was prepared according to the Promega protocol. RNA and primers were combined with reverse transcription mixture and incubated accordingly: Annealing at 25° C for 5 minutes; extension at 42° C for one hour and reverse transcriptase inactivation by incubation at 70° C for 15 minutes. The cDNA product was stored at −20° C prior to use.

Primers were designed to ensure that the entire coding region of the Ataxia– telangiectasia mutated (ATM) gene was amplified (Fig 1). Each set of primers were designed based on the wild type ATM (U82828.1, NCBI) sequence. Primers used in the experiment were designed as follows: primer length was between 18-25 base pairs; the primer melting temperature was calculated by A plasmid editor (ApE; http://biologylabs.utah.edu/jorgensen/wayned/ape/) software, with primers having a minimum melting temperature of 48°C; each primer was designed to have 40-60% GC content; and palindromic sequences within primers were avoided. Primer sequences are detailed in S1 Table. The use of a high fidelity, low error rate DNA polymerase enzyme in the PCR reactions was essential in order to minimise errors in amplicon extension and subsequent sequencing data. 1 µg cDNA samples were used as a template to generate full length high fidelity amplicons. The PCR reactions contained 0.5 μM forward primer, 0.5 μM reverse primer, 1 μl of cDNA sample, 1x Q5 PCR high fidelity Master Mix (NEB, MA, USA), with PCR quality water making the balance to 25 μl. PCR conditions for the different primer pairs are defined in S2 Table. PCR products were cleaned in 50 μl of elution buffer using the Ultra clean PCR clean up kit (Mo Bio, CA, USA). Confirmation of PCR product size was performed by 1 % agarose gel electrophoresis. Sanger sequencing was performed by the Australian Genome Research Facility in Brisbane and aligned with NCBI sequences of the ATM gene.

**Fig 1.**
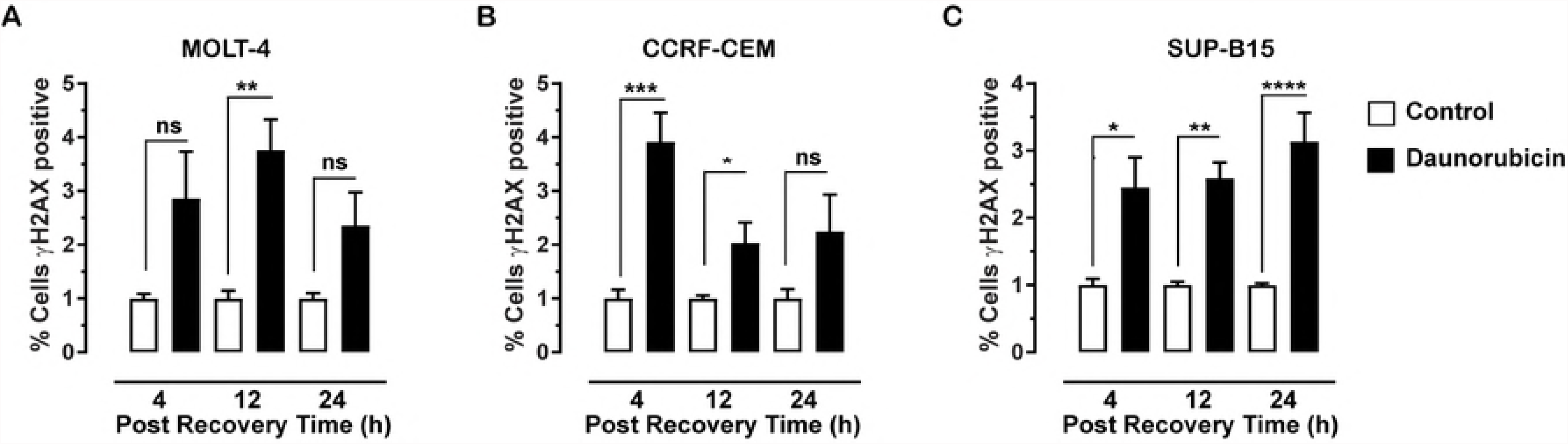
Schematic structure of the ATM gene and protein. **(A)** ATM cDNA diagram and primer positions. (**B)** ATM exon diagram and primer position. The red arrows indicate exons. The yellow arrows indicate size of cDNA sequence. The green arrows indicate primer position. Both **(A)** and **(B)** diagrams were created using CLC Genomics Workbench (CA, USA). **(C)** Schematic diagram of the regions of the ATM protein kinase. The FAT domain is autophosphorylated, PI3K is the kinase domain and FATC domain interacts with Tip60 protein to activate ATM.

## Data analysis

Data is presented as mean ± standard error of the mean (SEM) and is analysed by one-way ANOVA followed by the Tukey’s post-hoc test using GraphPad Prism 7 software. P < 0.05 is considered statistically significant.

## Results

### Effects of daunorubicin on cell viability

Daunorubicin treatment causes many different types of toxic lesions. The MTT assay was utilised to assess changes in the number of the three ALL cell lines used in this study. Each of the cell lines displayed a different pattern of sensitivity to daunorubicin. Daunorubicin toxicity was observed in MOLT-4 cells after 4 hours in recovery media, the earliest time examined (Fig 2A). The level of reduction in cell density was about 50% compared to the control after 4 hours (0.56 ± 0.05, P = 0.0018), and remained at these levels after 12 hours (0.54 ± 0.04, P = 0.0011) and (0.57 ± 0.02, P = 0.014) 24 hours. CCRF-CEM cells exposed to daunorubicin (Fig 2B) did not show significant reduction in cell density until 12 hours in the recovery media (0.48 ± 0.07, P = 0.0002) compared to the control. The significant reduction in the cell density level remained after 24 hours recovery (0.45 ± 0.07, P < 0.0001). Treatment of SUP-B15 cells (Fig 2C) with daunorubicin resulted in a biphasic response. Initially after 4 hours in recovery media there was a significant decrease in cell density compared to the control (0.61 ± 0.07, P = 0.006). When the cells were incubated in 12 hours recovery media, the decrease in cell density was comparable to the control (1.08 ± 0.07) and not significantly reduced at 24 hours (0.76 ± 0.06; P = 0.945).

**Fig 2.**
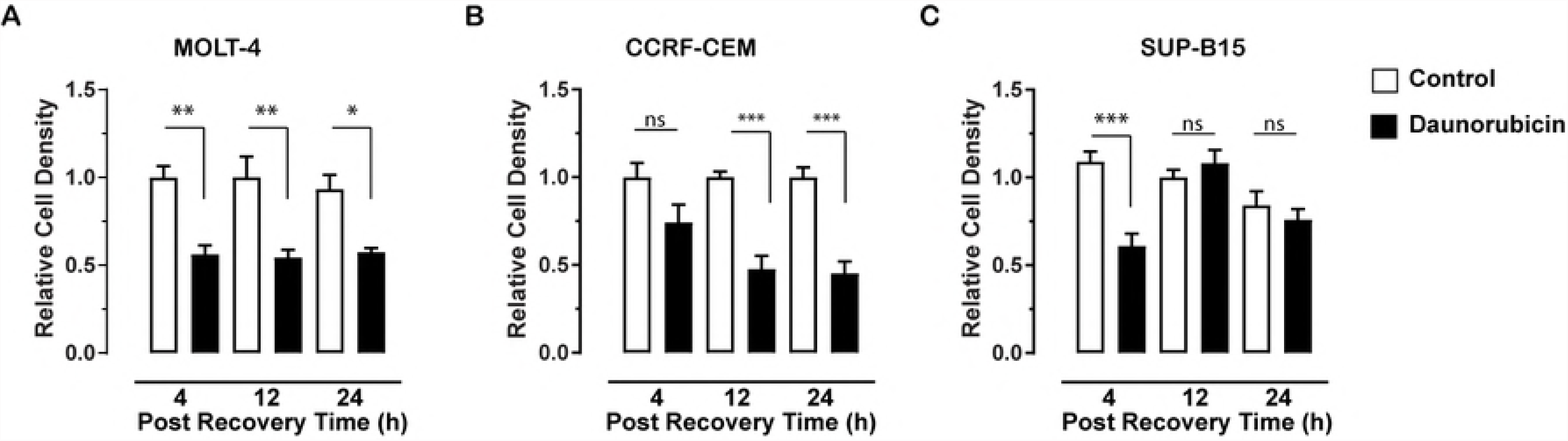
Effect of daunorubicin on cell viability. Relative cell density following treatment with 10 μM daunorubicin for 4 hours and followed by 4, 12 and 24 hours (post recovery time) of recovery media for (A) MOLT-4, (B) CCRF-CEM and (C) SUP-B15 cells was determined using the MTT assay. Bars indicate mean of a total of six replicates ± SEM, from three independent experiments. Results were normalised to the control at each time point. * P < 0.05, ** P < 0.01; *** P < 0.001 one-way ANOVA followed by Tukey’s post-hoc test.

### Effects of daunorubicin on production of damaging reactive oxygen species

A mechanism of action of daunorubicin is to generate reactive oxygen species (ROS), which causes DNA damage, particularly it will induce DSBs [16]. To assess the changes in ROS production over time following treatment with daunorubicin, the total ROS assay was performed (Fig 3 and S1 Fig). Since daunorubicin can stimulate the production of ROS via several processes, and cells have several mechanisms for quenching toxic ROS, the differences in ROS over time reflects not only cell sensitivity, but also long term daunorubicin effectiveness. Treatment of MOLT-4 cells (Fig 3A) with 10 μM daunorubicin resulted in a significant increase in ROS production after 4 hour recovery period (10.48 ± 0.03, P < 0.0001) compared to the control. Increase in ROS production reached a maximum after 12 hours recovery (127.7 ± 2.47, P < 0.0001) and declined significantly after 24 hours (24.90 ± 0.40, P < 0.0001). CCRF-CEM cells exposed to daunorubicin (Fig 3B), displayed a similar pattern of ROS production. There was a significant increase in ROS production after 4 hours recovery period (47.33 ± 0.77, P < 0.0001) compared to the control. This increase in ROS production reached a maximum after 12 hours recovery (133.1 ± 1.95, P < 0.0001), then decreased after 24 hours recovery (16.0 ± 0.21, P < 0.0001). SUP-B15 cells (Fig 3C) when treated with daunorubicin did not appear to be as sensitive to the production of ROS compared to MOLT and CCRF-CEM cells. SUP-B15 cells showed a significant increase in ROS production after 4 hours recovery period (23.09 ± 3.11, P < 0.0001) compared to the control. The amount of ROS produce was comparable following 12 hours incubation in the recovery media (24.75 ± 2.79; P < 0.0001) before declining after 24 hours recovery (9.91 ± 1.46 P = 0.0172).

**Fig 3.**
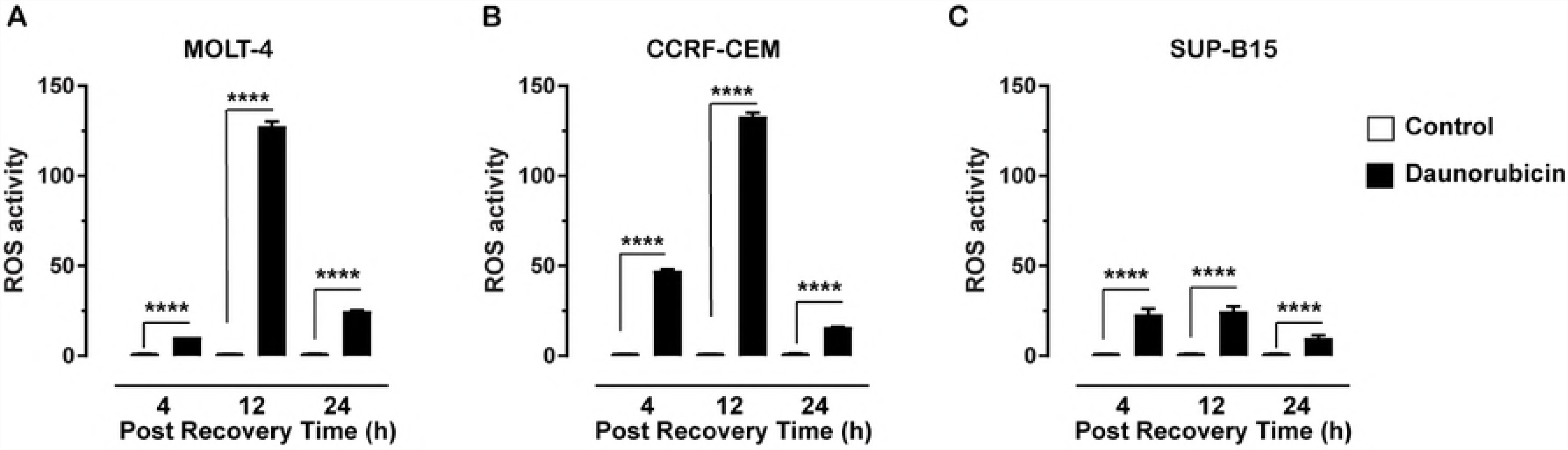
Total Reactive oxygen species (ROS) induced by daunorubicin. MOLT-4 (A), CCRF-CEM (B) and SUP-B15 (C) cells exposed to 4 h treatment with 10 µM daunorubicin, followed by 4, 12 and 24 hours (post recovery period) in recovery media. Graph indicates mean of total of six replicates ± SEM, from three independent experiments. Results were normalised to the control at each time point. * P < 0.05, ** P < 0.01; *** P < 0.001; **** P < 0.0001 one-way ANOVA followed by Tukey’s post-hoc test.

### Effects of daunorubicin on DNA double strand breaks

The lesions most detrimental to cell survival following treatment with daunorubicin, include the DSBs. The production and subsequent repair of these lesions was assessed over time by detecting γH2AX. This also gives an indication of how effective DNA repair processes are within T or B lymphoblast derived cells (Fig 4 and S2 Fig). After 4 hours recovery, there was no significant increase in γH2AX fluorescence intensity (2.81 ± 0.88; P = 0.127) compared to the control in MOLT-4 cells (Fig 4A). However, after 12 hours in recovery media, there was a significant increase in γH2AX expression (3.76 ± 0.57; P = 0.0065). When cells were incubated for 24 hours in the recovery media, γH2AX expression decreased and was comparable to control levels (2.35 ± 0.622; P = 0.423). When CCRF-CEM cells were allowed to recover for 4 hours (Fig 4B) following initial 4 hours treatment with daunorubicin, there was a significant increase in the number of cells that stained positive for γH2AX when compared to the control (3.91 ± 0.54; P = 0.0002). γH2AX expression decreased after 12 hours in recover media, comparable to control levels (2.37 ± 0.38; P = 0.471). Levels of γH2AX remained at this level after 24 hours recovery (2.24 ± 0.69; P = 0.277). When SUP-B15 cells (Fig 4C) were incubated for 4 hours in recovery media, there was a significant increase in the percentage of cells expressing γH2AX when incubated in the recovery media for 4 hours (2.45 ± 0.45; P = 0.0148). Expression levels remained elevated after 12 hours (2.59 ± 0.23; P = 0.0024) and 24 hours incubation in the recovery media (3.13 ± 0.43; P < 0.0001).

**Fig 4.**
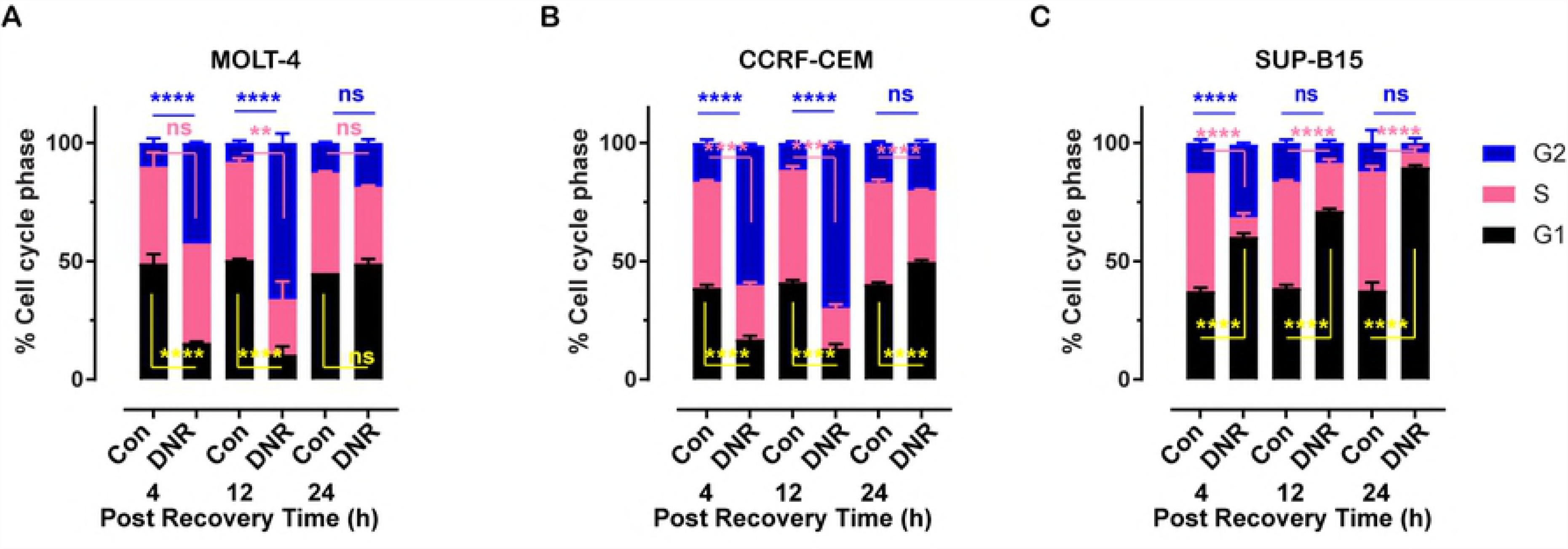
Effect of daunorubicin on DSB. Measurement of DSB by γH2AX fluorescence for MOLT-4 (A), CCRF-CEM (B) and SUP-B15 (C) cells following 4 h treatment with 10 µM daunorubicin, followed by 4, 12 and 24 hours (post recovery period) in recovery media. Median intensities from (flow cytometry histograms of the raw data) were used to plot the bar diagram. Graph indicates mean of total of six replicates ± SEM, from three independent experiments. Results were normalised to the control at each time point. * P < 0.05, ** P < 0.01; *** P < 0.001 one-way ANOVA followed by Tukey’s post-hoc test.

### Effects of daunorubicin on cell cycle progression

Following the toxic effects of daunorubicin, the cell should respond by initiating cell cycle arrest to allow DNA repair processes to occur. Analysis of cell cycle stages can be assessed using propidium iodide staining, to determine the impact of daunorubicin on cell cycle progression or arrest at different times after treatment (Fig 5). As shown in Fig 4A, cell cycle profiles for MOLT-4 after 4 hours in recovery media (15.5:42:42.5; G1:S:G2/M) there was an increase in the proportion of cells in G2/M phase of the cell cycle when compared to control (48:41:11), with a subsequent decrease in G1 phases. The proportion of cells continued to accumulate in the G2/M phase after 12 hours recovery (10.5:23.5:66) with further reduction in G1 phase, but after 12 hours there was also a reduction in cells in S phase. After 24 hours recovery time, the cell cycle profile returned to normal levels (49:32.5:18.5). Similarly, the cell cycle profiles for CCRF-CEM cells (Fig 5B) showed a dramatic increase in the proportional of cells in G2/M phases of cell cycle when compared to control. The proportion of cells in the G2/M phase continued to increase after 12 hours recovery. The accumulation of cells in the G2/M phase resulted in a reduction of cells in the G1 and S phase of cell cycle. After 24 hours recovery time, the cell cycle profile returned to normal levels. In contrast, the cell cycle profiles for SUP-B15 cells (Fig 5C) showed an increase in the proportion of cells in G1 phase and G2/M phase of cell cycle when compared to control after 4 hours of incubation in recovery media. The proportion of cells in G1 phase was further increased when the cells were incubated after 12 hours, further increasing after 24 hours recovery. Cells accumulating in the G1 phase resulted in a reduction of cells in the G2/M and S phase of cell cycle.

**Fig 5.**
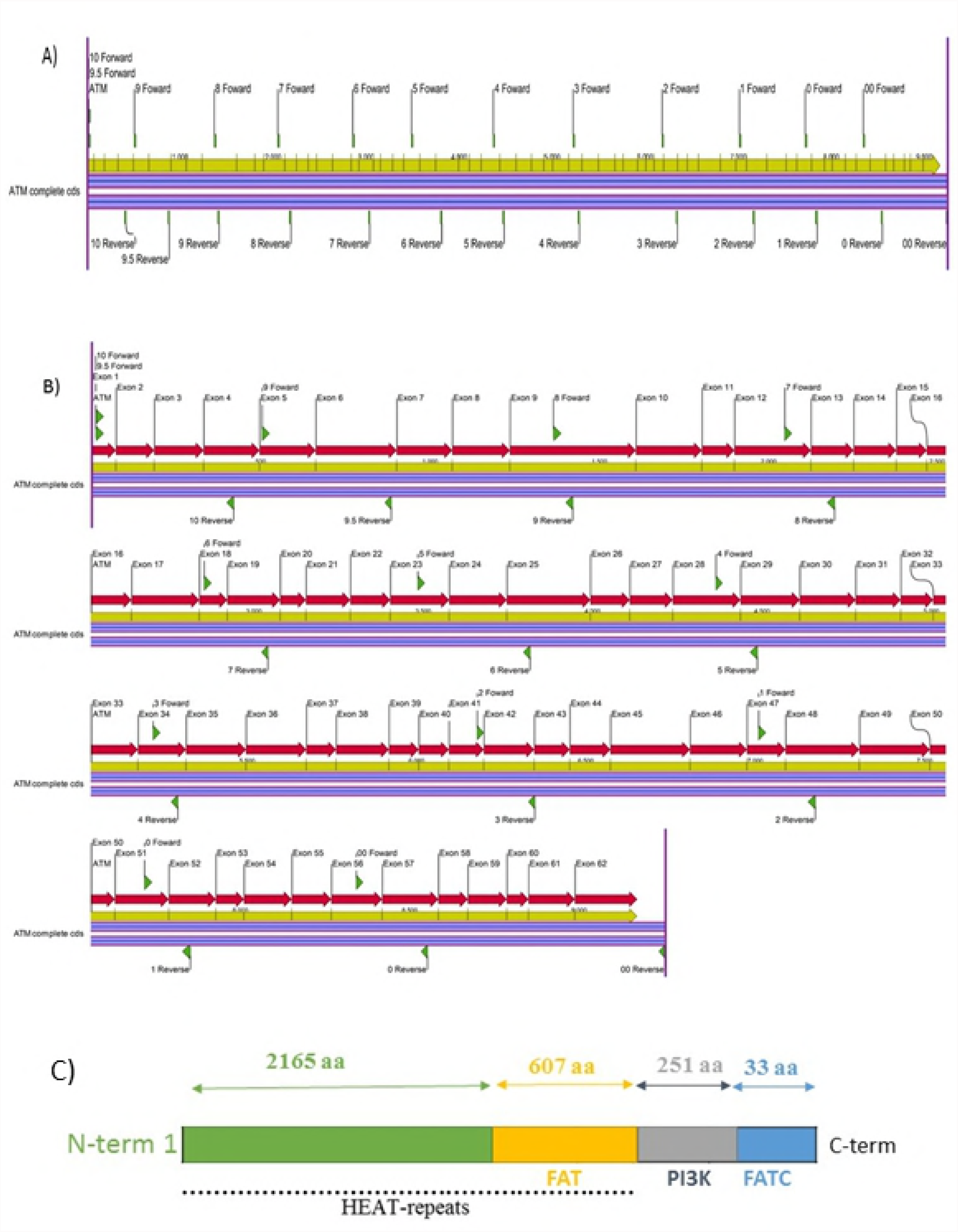
Daunorubicin alters cell cycle profiles. Cell cycle profiles for MOLT-4 (A), CCRF-CEM (B) and SUP-B15 (C) cells following 4 h treatment with 10 µM daunorubicin followed by 4 hour, 12 hour and 24 hour in recovery media. Graph indicates mean of total of six replicates ± SEM, from three independent experiments. * Results were normalised to the control at each time point. * P < 0.05, ** P < 0.01; *** P < 0.001; ns = non-significant one-way ANOVA followed by Tukey’s post-hoc test. Black G1 phase, Pink S phase and Blue G2/M phase.

### Expression of p53 and SOD2

DSB activates ATM [17] and the differences in cell cycle recovery may be due to mutations in ATM. To determine if functional mutations in ATM were affecting how cells handle chemotherapy treatment, the full ATM sequence in T-lymphoblast derived CCRF-CEM and MOLT-4 cells and B-lymphoblast derived SUP-B15 cells were explored through Sanger sequencing. The coding region was fully amplified using a selection of primer pairs (Fig 1) and the products were sequenced. This was done for each of the cell lines and no mutations found in the full ATM coding region.

Following detection of DSB, activation of p53 initiates several downstream processes, including activation of SOD2 to quench ROS, and maintain cell survival. Analysis of p53 in the three ALL cell lines following treatment, indicated a lack of p53 phosphorylation at Ser15 in SUP-B15 cells. Whereas the phosphorylation of p53 in both CCRF-CEM and MOLT4 cell lines was observed (Fig 6A). SOD2 was unchanged in both MOLT-4 and CCRF-CEM and increased in SUP-B15 cells (Fig 6B), although the later effect could be due to reduced ROS production seen in SUP-B15 cells (Fig 3C). This dysfunction of p53 would also have an impact on cell cycle regulators, including p21 and p27, which are both down regulated in SUP-B15 cells following treatment with daunorubicin (Table 1).

**Fig 6.**
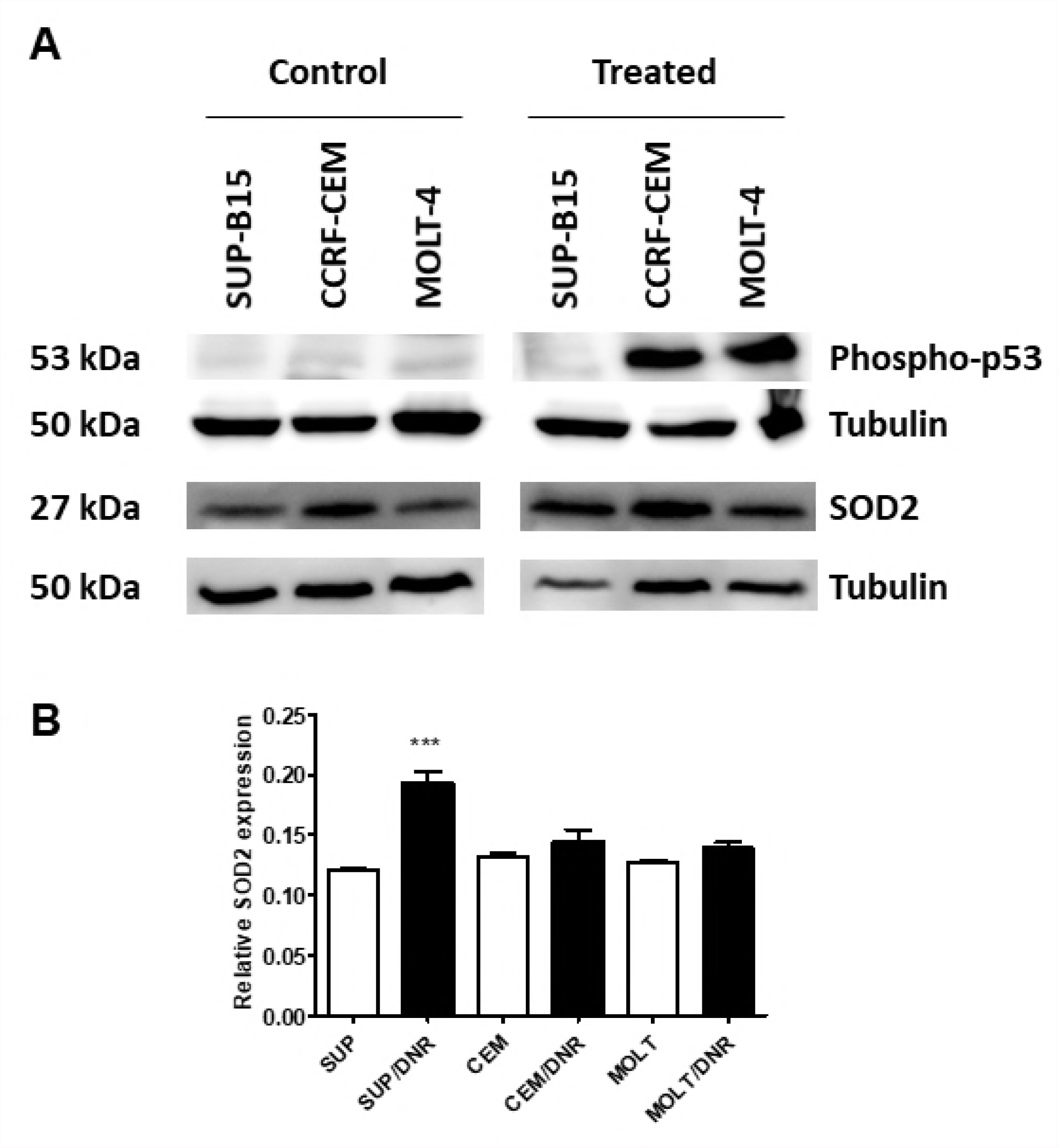
Western analysis of p53 and SOD2. **(A)** Proteins from SUP-B15, CCRF-CEM and MOLT-4 cell lines treated with 10 µM daunorubicin for 4 h or untreated (control) were probed with anti-SOD2 or anti-phospho-p53 at Ser15. Separate gels from the same cell preparations were analysed for tubulin expression. **(B)** Evaluation of MnSOD2 expression. Tubulin expression was analysed as a loading control, results are expressed as ratio of SOD2 to tubulin intensity. Bars indicate mean of at least four replicates ± SEM, from two independent experiments. *** P < 0.001 one-way ANOVA.

**Table 1.** Summary of results from the apoptosis array showing the effect daunorubicin on p53, p21 and p27 in MOL-4, CCRF CEM and SUP-B15 cell lines.

| Protein | MOLT-4 |  | CCRF-CEM |  | SUP-B15 |  |
| --- | --- | --- | --- | --- | --- | --- |
|  | DNR | Recovery | DNR | Recovery | DNR | Recovery |
|  | 4 hours | Media<br>12 hours | 4 hours | Media<br>12 hours | 4 hours | Media<br>12 hours |
| **p53** | ↑ | — | ↑ | — | ↓ | ↓ |
| **p21** | ↑ | — | ↑ | — | ↓ | ↓ |
| **p27** | ↑ | - | ↑ | ↑ | ↓ | - |

## Discussion

Assessment of the impact of daunorubicin on the selected leukaemic cell lines revealed very different cellular responses and sensitivity to daunorubicin. The ALL cell lines, MOLT-4 and CCRF-CEM which are derived from acute T-lymphoblastic leukemia, displayed a cytotoxic response, with DNA-DSB and cell cycle arrest to allow subsequent DNA repair to occur. However, the ALL cell line, SUP-B15 derived from an acute B-lymphoblastic leukaemia, displayed a different pattern of response. SUP-B15 cells showed no signs of DNA-DSB repair, and cells accumulating in G1 phase, as opposed to G2/M in the other cell lines. Regulation of cell cycle progression and DNA repair through activation of p53, p21 and p27, was reduced differing from the observed in MOLT4 and CCRF-CEM cells.

All cell lines in this study responded to the cytotoxic effects of daunorubicin with a reduction in cell number which supports previous studies [13, 18, 19]. MOLT-4 cells appeared to be the most sensitive, with a persistent decrease in cell number observed as early as 4 hours post treatment. CCRF-CEM cells did not succumb to the cytotoxicity until 12 hours post treatment, levels at this time were comparable to MOLT-4, and remained at this level for the duration of the study time. However, SUP-B15 cells displayed a different pattern of response and appeared less sensitive to the drug especially after 12 and 24 hours. These differences suggest that daunorubicin induced more DNA damage in both MOLT-4 and CCRF-CEM cells compared to SUP-B15 cells. The concentration of daunorubicin (10 μM) used in this study was shown to be effective in Jurkat T lymphoma and HL-60 promyelocytic leukaemia cell line [20] and sarcoplasmic reticulum cardiac cells [21].

The effectiveness of a chemotherapeutic agent is dependent on several factors; concentration, exposure time, doubling time of the cell line, state of DNA Damage Response (DDR) mechanisms and type of damage induced. Most of these characteristics are primarily determined by the genetics of the cell line. Many chemotherapeutic agents, such as daunorubicin, elicit their damage by disrupting or targeting the replication of DNA during the S phase of the cell cycle. Cells that complete more cell cycles (have shorter doubling times) are therefore more vulnerable to the chemotherapeutic agent when compared to a cell line that completes fewer cell cycles (longer doubling time). The CCRF-CEM cells have a doubling time ranging from 20 to 30 hours [22], therefore the CCRF-CEM cells would have at least one cell cycle to identify and repair the damage induced by the chemotherapeutic tested. MOLT-4 cells have a doubling time of approximately 22-24 hours [23-25]. So again, MOLT-4 would have one cell cycle to identify and repair the damage induced by the chemotherapeutic agents. SUP-B15 cells would have only entered the second cell cycle during the observed treatment time as the doubling time of SUP-B15 cells is approximately 46 hours, with reports ranging from 35 to 60 hours [26, 27]. This may explain the observed results as half of SUP-B15 cells have been affected, while MOLT-4 and CCRF-CEM the majority of cells have been damaged.

Free radical formation and oxidative stress play an important role in the cytotoxicity of daunorubicin as ROS may serve as an intracellular signal of apoptotic events [28]. Daunorubicin induced ROS in the three leukaemic cells lines with variations over the time period. The T-lymphoblast cell lines, CCRF-CEM and MOLT-4 produced ROS at high levels from 4 to 12 hours before declining sharply over the next 12 hours. However, the B lymphoblast SUP-B15 cells consistently expressed significant but relatively lower ROS levels throughout the experimental period. Daunorubicin induces cytotoxicity in the cells by the generation of ROS and enhanced G2/M phase cell cycle [29]. Increases in the level of ROS is regulated by several signal networks, Antioxidants such as superoxide dismutase (SOD2), catalase (CAT) and glutathione peroxidase 1 (GPx) play a central role [30]. The overall effect of the antioxidant system depends on the intracellular balance between these antioxidant enzymes rather than a single component. In the antioxidant enzyme system, SOD2 catalyses the dismutation of superoxide radicals to H_2_O_2_. p53-dependent up-regulation of SOD2 (or MnSOD) and glutathione peroxidase 1 (GPx) is important in human lymphoblasts [31].

Although ATM was expressed in all three cell lines and contained no mutations in the coding regions, p53 was phosphorylated in MOLT-4 and CCRF-CEM but not in SUP-B15 after daunorubicin treatment. Hence, p53 in SUP-B15 is more likely to be mutated. Leukaemia cells often possess mutations in p53 gene [32]. P53 is an essential tumour suppressor gene, that also regulates SOD2 gene expression [33]. Thus in SUP-B15 cells where p53 is not activated, the antioxidant effects of SOD2 will not modulate ROS levels during the daunorubicin treatment [34]. However, p53-mediated production of ROS can be cell type and species dependent [31].

To determine the impact of daunorubicin on the DSBs in the selected ALL cell lines, and the subsequent repair, we analysed the gamma H2AX levels. The histone H2AX plays a key role in DNA-DSB repair by rapidly phosphorylating serine residues to form γH2AX foci near the DSBs [35]. Treatment of all cell lines with daunorubicin resulted in an increase in γH2AX staining within 4 hours indicating this is an appropriate treatment time and supporting previous studies that show daunorubicin causes DSBs [36, 37]. To follow the dynamics of DSB repair, cells were allowed to recover over 24 hours when there were still significant levels of DSBs detected. This indicates that even though repair was occurring, not all DBSs induced were repaired during this time. Notably, SUP-B15 cell line showed more pronounced DSBs at 12 and 24 hours recovery time compared to MOLT-4 and CCRF-CEM cell lines, suggesting that different DDR mechanisms or pathways were involved. DSB repair in CCRF-CEM cells appeared the most robust, with repair occurring after 4 hours in recovery and total repair after 24 hours. MOLT-4 showed that the DSB repair took place after 12 hours in recovery. This indicates that the 24 hours recovery time is enough for some repair of daunorubicin induced DSB in MOLT-4 and CCRF-CEM cells. On the other hand, even after 24 hours recovery, the level of DSBs in SUP-B15 cells was significantly higher indicating incomplete DSB repair. Most cells have an intrinsic repair process in response to any DNA damage, including those induced by daunorubicin. The effectiveness of the DDR will be cell line dependent, with some cell lines having mutations in the key signalling intermediary p53 [38, 39] as seen in SUP-B15 in this study. Dependent on the DNA damage that has been induced, single stranded breaks or DSB, different DDR pathways will be signalled. Daunorubicin induced DNA-DSB primarily utilise the HR and NHEJ repair pathways [40]. Mutations to important elements of the HR or NHEJ, can compromise the DDR mechanisms, resulting in less damage being identified and appropriate cellular responses stimulated. Increased expression of such enzymes leads to increased repair following exposure to chemotherapies inducing DSB, including daunorubicin, and this is a key mechanism in the ever growing problem of chemoresistance to therapy [41, 42]. The difference in potency of the chemotherapy in the three cell lines could be due to the difference in molecular profiles between the three cell lines, and one pertinent example is p53 status.

Along with reduction of detectable γ-H2AX, and thus DSBs, during the recovery time, DNA-PK, ATM and ATR also initiate cell cycle arrest. Exposing SUP-B15 cells to daunorubicin caused a progressive accumulation of cells in G1 phase while daunorubicin treatment of MOLT-4 and CCRF-CEM cells caused a profound accumulation of cells in G2/M phase, with a progressive reduction of cells in G1 and S phase of the cell cycle. However, after 24 hours recovery the cell cycle profile of MOLT-4 and CCRF-CEM cells was comparable to the control, suggesting the impact on cell cycle was no longer expressed. This is consistent with the previous observation in HL-60 cells (myeloid leukaemic cell line) where daunorubicin caused a marked G2/M accumulation after 24 hours of exposure to the drug [43]. The profound accumulation in G2/M has also been observed in CCRF-CEM cells treated with the anthracycline doxorubicin [44]. Doxorubicin also induced profound G2/M arrest in HCT-116 human colon carcinoma cells and was accompanied by activation of p53 and induction of p21 mRNA and protein expression [45].

The cell line dependent variations in various enzyme expression levels, particularly p53, p27 and p21, could be a factor in the differences in the degree of cell cycle arrest and subsequent DSB repair. In many cancer cells, loss of p53 is thought to be a predictor of failure to respond to chemotherapy and radiotherapy [46]. Treatment of MOLT-4 and CCRF-CEM with daunorubicin resulted in increased the level of p53, p27 and p21, corresponding with increased levels of DNA DSBs and cell cycle arrest. In contrast, SUP-B15 cells showed decreases in the levels of p53, p27 and p21. Several studies have provided compelling evidence relating the mutations in p53 with chemoresistance to various cytotoxic drugs [38, 47-50].

Following DNA damage, phosphorylation of p53 at Ser15 by ATM or other kinases inhibits the binding of MDM2 to p53 and this leads to increased activity of p53. Anthracycline mediated cell cycle arrest may take place at G1 or G2 checkpoints and this can be mediated through the multifunctional transcription factor p53 [51, 52]. p53 induces p21 (CDK inhibitor) expression and therefore, inactivation of p53 will also result in decreasing levels of p21 [53]. p27 is also a member of CIP/KIP and controls cell progression from G1 to S phase by mediating G1 arrest through inhibiting cyclin/CDK complex activities [54]. Normally, p53 may boost chemosensitivity through enhancing p21-mediated growth arrest and DNA repair [55]. p53 has two functional roles, it can induce cell cycle arrest, through the transactivation of the p21 or induction of apoptosis [56].

## Conclusions

In summary, the study provides additional insight into the mechanism of action of daunorubicin on DNA DSB formation, DNA repair and the cell cycle arrest in acute lymphoblastic leukaemia cell lines. The delay in DSB repair and lower sensitivity to daunorubicin in the B lymphoblastic SUP-B15 cells is likely to involve loss of p53 function amongst other factors. These factors may contribute in inhibiting or affecting DNA repair pathways. As p53, p21 and p27 phospho-kinase proteins play essential roles in tumour progression and clinical outcomes in acute lymphoblastic leukaemia, the presence and activity regulatory proteins should be taken into consideration in devising personalized treatment regimens.

## Acknowledgements

The authors would like to thank La Trobe Institute of Molecular Sciences (LIMS) and La Trobe University for supporting the research. Hussain Al-Aamri would like to thank the Ministry of Higher Education, Government of Oman for providing a Postgraduate Research Scholarship.

## Supporting Information

**S1 Fig. Examples of histograms used for the ROS assay.** Illustrative histograms obtained from (A) MOLT-4, (B) CCRF-CEM, and (C) SUP-B15 cell lines.

**S2 Fig. Examples of histograms used for the gamma H2AX assay.** Illustrative histograms obtained from (A) MOLT-4, (B) CCRF-CEM, and (C) SUP-B15 cell lines

**S1 Table. Primers used to amplify the ATM cDNA.**

**S2 Table. PCR conditions for the different primer sets to amplify ATM.**

